# Role of Udd protein and heterochromatin in transcriptional selection of individual rRNA genes in the *Drosophila* germline

**DOI:** 10.1101/2020.10.21.349613

**Authors:** Elena A. Fefelova, Irina M. Pleshakova, Sergei A. Pirogov, Elena A. Mikhaleva, Valentin A. Poltorachenko, Roman S. Blokh, Yuri A. Abramov, Daniil D. Romashin, Vladimir A. Gvozdev, Mikhail S. Klenov

## Abstract

Eukaryotic genomes contain hundreds of nearly identical rRNA genes, many of which are transcriptionally silent. However, the mechanisms of selective regulation of individual rDNA units remain poorly understood. In *Drosophila melanogaster*, rDNA repeats containing insertions of R1/R2 retrotransposons within the 28S rRNA sequence undergo inactivation. Here we found that rRNA genes with insertions are specifically enriched with H3K9me3 and HP1a repressive marks, but disruption of heterochromatin components only slightly affects their silencing. Intriguingly, the loss of Udd (Under-developed) protein interacting with Pol I transcription initiation complex, causes an upregulation of R2-inserted rDNA copies in germ cells by two orders of magnitude that is accompanied by the reduction of heterochromatin marks. Thus, for the first time we revealed a factor required for distinguishing between active and silent rDNA units to such a large extent. To clarify a relationship between the rDNA transcriptional status and heterochromatin establishment, we showed that inhibition of transcription by actinomycin D increases the level of H3K9me3 mark erasing the epigenetic differences between inserted and uninserted rRNA genes. Altogether, we suggest that Udd coupled with Pol I transcription initiation machinery defines activation or silencing of individual rDNA units, whereas their transcription level consequently dictates their chromatin state.

## Introduction

Regulation of ribosomal DNA (rDNA) transcription is a fine-tuned mechanism that defines the level of protein synthesis and controls cell growth and differentiation (1,2). Production of rRNA by RNA polymerase I (Pol I) occurs in the nucleolus accounting for up to 60% of transcriptional activity in the metabolically active eukaryotic cell (3,4). Eukaryotic rDNA is organized in clusters (also known as the nucleolus organizer regions (NORs)), generally consisting of hundreds of tandemly repeated rRNA genes separated by intergenic spacer regions (IGS) (5). Each gene expresses a pre-rRNA transcript harboring an external transcribed spacer (ETS) followed by the sequences of 18S, 5.8S and 28S rRNAs, which are interspaced by internal transcribed spacers (ITSs). Both ETS and ITSs are eliminated during nuclease processing steps, leading to formation of mature rRNAs.

Numerous genetic, microscopic and biochemical studies performed on a variety of objects have demonstrated that only a part of rDNA units is transcriptionally active at any time, while the rest, or even the majority of rDNA repeats, are in an inactivated state (6-14). In organisms whose genomes contain several rDNA clusters, silencing can occur at the level of entire NORs. This phenomenon, generally called nucleolar dominance or allelic inactivation of rDNA loci, is well described in plants, mammals and the fruit fly (15-18). Moreover, within the active NORs, some rRNA genes can be silent (19,20). For example, the clustering of both active hypomethylated and repressed hypermethylated rDNA units was revealed in human fibroblasts and carcinoma cells (20). Repression of a fraction of rRNA genes can perform various functions. First, this phenomenon can be useful to maintain an optimal level of ribosome production regardless of the rDNA copy number in the genome (21). Secondly, maintaining a subset of rRNA genes in an inactive heterochromatic state is thought to be important for ensuring nucleolar structure as well as preventing recombination between rDNA repeats (22-24). Besides, it is necessary to repress transcription of defective rDNA units, which are abundant in eukaryotic genomes. For instance, approximately 20–30% of rDNA repeats in human cell lines are palindromic and noncanonical (20,25). Repressive histone modifications, DNA methylation, and nucleosome remodeling etc. have been revealed to be among mechanisms responsible for rDNA silencing (see (26-29) for reviews). However, it remains generally unknown how cellular machineries select particular rRNA genes to establish their activation or repression.

*Drosophila* provides an attractive model for studying the transcriptional regulation at the level of individual rDNA units because some rRNA genes in this organism contain insertions of non-LTR retrotransposons called R1 and R2 (see (30,31) for reviews). In *D. melanogaster*, rDNA clusters located in the X and Y chromosomes usually harbor between 200-250 rDNA units (32) wherein more than half of them are inserted. One study showed that R1 elements interrupt from between 15% to 67% of rRNA genes, and R2 up to ∼ 30% depending on *Drosophila* strain (33). The R2 element is highly conservative being present in the genomes of various groups of animals, but not found in mammals (30,34). R2 retrotransposons insert into a strictly defined site in the rDNA sequence located 2651 bp after the beginning of the 28S rRNA, while R1 elements are usually integrated 74 bp downstream of the R2 insertion site (30). R2 elements are of particular interest because they are present exclusively at the rDNA locus, whereas R1 insertions have also been found in other regions of the genome (35-38). Along with unusually high specificity of integration, another unique feature of R2 elements is that they do not have their own promoters and therefore can be transcribed only as a part of the pre-rRNA transcripts (39-42). Hence, R2 sequences can be considered as markers that allow detection of the expression of corresponding rDNA units. Full-length R2 insertions encode a self-cleaving ribozyme, which releases the 5’-end of the R2 transcript from the upstream 28S rRNA sequence during the autocatalytic reaction (Figure 1A), whereas the mechanism of 3’-end formation of R2 RNA remains unclear (30,42). The protein product encoded by this element, namely R2 protein, has several domains, including DNA- and RNA-binding motifs, a reverse transcriptase (RT) and an endonuclease (EN). The EN domain harbors an active site similar to that of some restriction enzymes, defining the specificity of R2 integration into 28S rDNA. The RT domain performs reverse transcription coupled with the integration of a new R2 copy into the genome (30,43-46).

**Figure 1.**
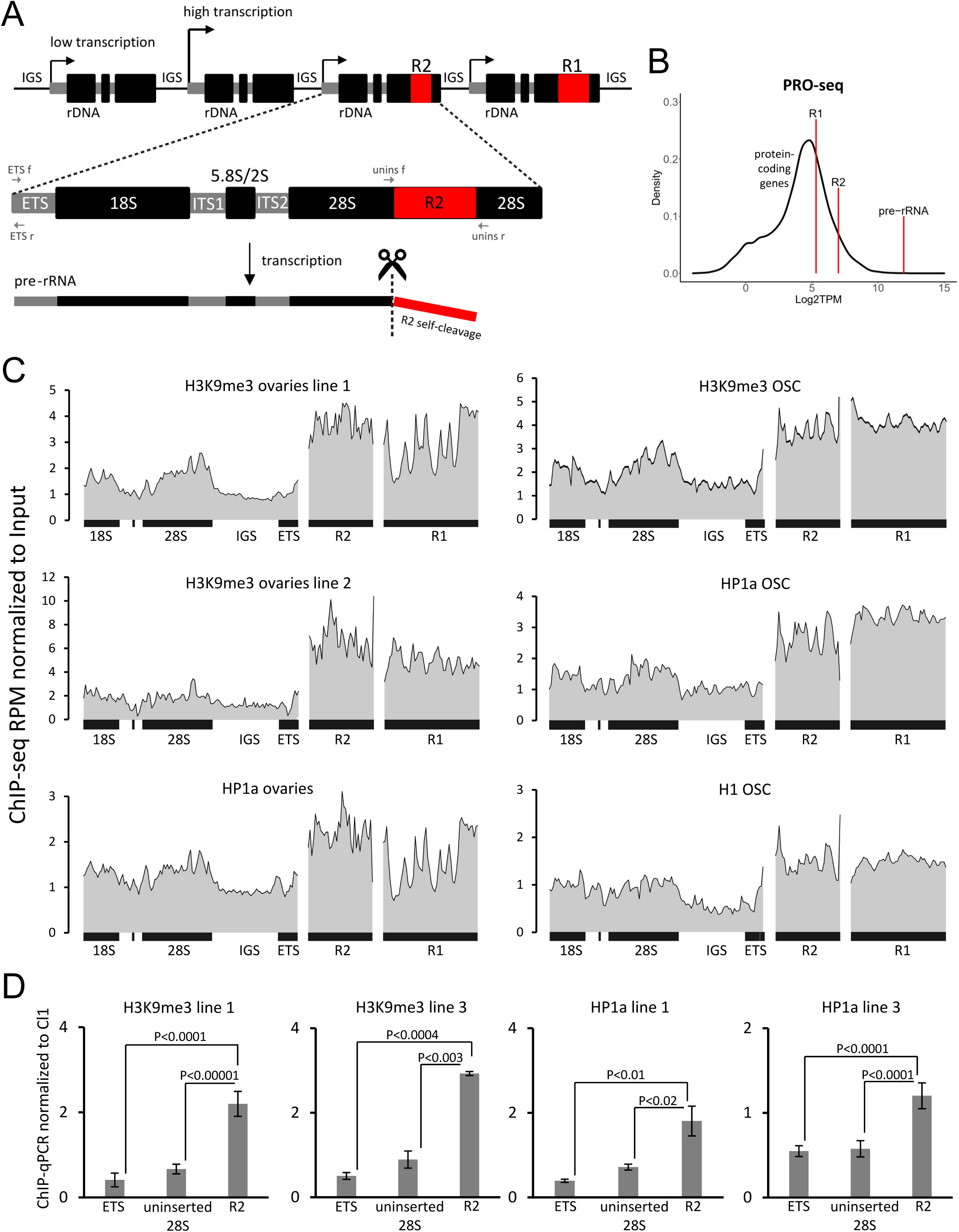
Inserted rDNA units are enriched in heterochromatin marks compared to uninserted ones. **(A)** Scheme of the rDNA array fragment in *D. melanogaster*. The rDNA cluster consists usually of 200-250 rDNA units separated by intergenic spacers (IGS). Each rRNA gene is about 8 kb long and encodes 18S, 28S and 5.8S rRNAs, as well as a small 2S molecule. Some rRNA genes contain insertions of R1 and R2 elements in the 28S sequences. R2 transposons are transcribed only as a part of pre-rRNA molecules. A self-cleaving ribozyme releases the 5’-end of the R2 transcript from the upstream 28S rRNA sequence. Location of primers used for detection of uninserted rDNA genes and ETS region is indicated by arrows. (**B)** Input-normalized levels of transcription of single rDNA repeat, R1 and R2 elements according to ovarian PRO-seq. The distribution of transcription levels (log2 TPM) of single-copy protein-coding genes is shown by black line. (**C)** Density profiles of ChIP-seq reads on rDNA unit, R2 and R1 elements in ovaries and OSC cells. RPMs are normalized to the corresponding Input DNA in 100 bp windows. The genotypes of the analyzed *Drosophila* lines are indicated in the Materials and Methods. (**D)** ChIP-qPCR analysis of H3K9me3 and HP1a levels in ovaries of two *D. melanogaster* lines. The enrichment levels of uninserted 28S sequence, R2 insertions and the beginning of ETS, which corresponds to promoter regions of both inserted and uninserted rDNA units, are normalized to the heterochromatic cluster-1 (Cl1, 42AB). Mean +/−s.d. and p-values based on two-sided Student’s t-test are indicated.

Given that R2 elements hijack the most potent transcription machinery in eukaryotic cells, namely Pol I, they are likely to be under especially strong control from cellular silencing mechanisms. With complete derepression, one would expect an accumulation of their transcripts comparable to rRNA abundance. However, it has been known for several decades that rRNA genes with insertions are usually transcribed at levels hundreds or thousands of times lower than uninserted rDNAs (47-49), though in one work only about a 10-fold difference was observed (50). Electron microscopy using the “Miller spreading technique” (6) showed that the low level of R2 transcripts is mainly due to the transcriptional repression of the entire rDNA units, which contain insertions (51,52). However, run-on analysis indicated that transcription of inserted rDNA repeat can be initiated, but is often terminated within the insertion sequence (50). Furthermore, it was suggested that most uninserted rRNA genes are also silenced: psoralen cross-linking assay revealed that only less than 10% of the total rDNA units are actively transcribed (50). The mechanism of inserted rDNA recognition and repression remains obscure, though some assumptions have been made in a number of reports (40,53,54). In particular, it was supposed to engage small RNAs and local formation of heterochromatin (30), i.e. the pathways that are known to determine silencing of conventional transposable elements (TEs) in the *Drosophila* genome (55-57). An upregulation of R1 and R2 (usually several times relative to controls) was observed in some experimental systems due to the impairment of Ago2-siRNA and Piwi-piRNA pathways (58-61), H1 histone (62), lamin (63), the architectural protein CTCF (53), and the nucleolar protein Nopp140 (64). An analysis of the proportion and interposition of inserted and normal rRNA genes within clusters suggested a “transcriptional domains” model according to which the region of the rDNA cluster lacking R1/R2 insertions is selected to be transcriptionally active, while the rest of the locus undergoes heterochromatinization (40,65). However, differences in chromatin density and histone modifications between inserted and uninserted rDNA units have not been observed in work on embryonic cells (50). Nevertheless, another report found the colocalization of R1 and R2 elements with condensed chromatin in the nucleolus of salivary gland cells (38). Overall, the role of chromatin context in the repression of inserted rDNA units, as well as in the regulation of *Drosophila* rDNA transcription in general remains controversial and under-investigated.

Here, we study the silencing of rRNA genes with TE insertions in *D. melanogaster* ovaries. Ovarian germ cells are characterized by an extremely intense rRNA synthesis required for the oocyte development (66,67), as well as by a high activity of various TEs, which tend to transpose within the genome of gamete precursor cells (57). We show that the chromatin of inserted rDNA units in ovaries is enriched with H3K9me3 and HP1a repressive chromatin marks compared to uninserted rDNA units, which for the first time demonstrates the existence of differences in chromatin states between individual rDNA repeats in *Drosophila*. However, disruption of H3K9 methylation or HP1a leads to a weak upregulation of inserted rRNA genes, whereas the loss of Under-developed (Udd) protein, previously identified as a component of SL1-like complex responsible for Pol I transcription initiation (68) causes a drastic derepression of rDNA units with insertions in the germ cells. The loss of Udd was also accompanied by the reduction of heterochromatin marks associated with rDNA chromatin. On the contrary, inhibition of transcription by actinomycin D treatment increased the level of H3K9me3 in the chromatin of uninserted rRNA genes. We therefore hypothesize that Udd is responsible for rDNA “transcriptional selection” that normally defines the transcription level of rRNA genes, whereas heterochromatin marks are further recruited on silent or less effectively transcribed rDNA copies.

## Materials and methods

### *Drosophila* strains, maintenance, and crosses

*Drosophila melanogaster* stocks and crosses were maintained under standard conditions at 25°C or in some cases, as indicated below, at 18°C. *udd*^*1*^ and *udd*^*null*^ lines were kindly provided by M. Buszczak (68). For analysis of *udd* mutant phenotype, *udd*^*1*^*/udd*^*1*^ flies in comparison with *udd*^*1*^/+ control or *udd*^*1*^*/udd*^*null*^ flies in comparison with heterozygotes obtained in the same cross (mix of *udd*^*1*^/+ and *udd*^*null*^*/+*, designated in the text as *udd/+*) were used. Because of the increasing with age loss of germ cells in *udd*^*1*^*/udd*^*1*^ and *udd*^*1*^*/udd*^*null*^ ovaries, for most experiments the ovaries from 0-1day old females were isolated. For ChIP-qPCR analysis we used the wild type *Batumi* strain (designated as line 3) and *piwi*^*2*^/+;*piwi*^*Nt*^/+ flies (designated as line 1) exhibiting the wild-type ovary phenotype (69). This line was used as a model in some ChIP-qPCR experiments, since H3K9me3 and HP1a ChIP-seqs were previously made for this line (70).

The following UAS-RNAi stocks were obtained from Vienna Drosophila Resource Center (VDRC): egg RNAi (#101677, #109673), Su(var)3-9 RNAi (#39378, #101494), HP1a RNAi (#31994, #31995), white RNAi (# 30033). The resulted RT-PCR values were averaged based on three biological replicates for each of the two RNAi lines for egg, Su(var)3-9 and HP1a. Germline knockdowns (GKD) were induced by crossing these lines with *nos-GAL4* driver *P*{*UAS-Dcr-2*.*D*}*1, w*^*1118*^, *P{GAL4-nos*.*NGT}40* (#25751 Bloomington Drosophila Stock Center (BDSC)), providing germline-specific GAL4 expression under the control of the *nanos* (*nos*) gene promoter. HP1a and Su(var)3-9 GKD crosses were grown at 18°C, since maintaining at 25°C resulted in almost complete loss of germ cells.

Fly stocks for histone replacement system (71) were kindly provided by Robert J. Duronio. These stocks contain a transgenic histone cluster, in which codons for K9 lysine residues of the H3 histone genes are substituted with arginine (K9R), or a wild type transgenic histone cluster (HWT). Progeny lacking endogenous histone genes and containing HWT or K9R transgenes integrated on chromosome 3, was produced by crossing parents heterozygous for the *HisC* deletion on chromosome 2 and identified by YFP expression using UAS-GAL4 as described (71). For this, *ΔHisC, twi-GAL4/CyO* females were crossed with *ΔHisC, UAS-2xEYFP/CyO; HWT/HWT* or *K9R/K9R* males. Therefore YFP-positive progeny carried *HisC* deletion on both chromosomes 2. Then YFP-positive HWT and K9R larvae were manually selected and used for RT-qPCR analysis. The absence of H3K9me3 modification in K9R larvae was confirmed by immunostaining of polytene chromosomes of salivary gland cells with H3K9me3 antibodies (Upstate).

### Chromatin immunoprecipitation (ChIP)

ChIP was performed according to the published procedure (72) with some modifications. Formaldehyde cross-linking was performed immediately after manual ovary dissection, that was found to be important to obtain high enrichments. Chromatin was fragmented by sonication on Vibra-Cell (Sonics) with amplitude 15% during 35 pulses of 10s with 10s pause intervals. 0.6 μg of chromatin extract was taken for DNA Input and stored at -20°C. For IP, 2 μg of chromatin extract was diluted up to 500 μL in lysis buffer and pre-incubated overnight at 4°C in the presence of 50 μL Protein A-Sepharose suspension (Amersham Pharmacia, 50% w/v, hereinafter PAS). After removing PAS, samples were incubated at 4°C for 5 h with antibodies and overnight with the addition of PAS. Then, samples were centrifuged at 3000 x g for 1 min and the supernatant was discarded. Subsequent steps were performed as described (72). Chromatin was immunoprecipitated with the following antibodies: rabbit α-H3K9me3 (Upstate), rabbit α-HP1a (a gift from Sarah Elgin), rabbit α-H4K20me3 (Abcam, ab9053), α-H3K27me3 (Upstate), guinea pig α-Udd (a gift from Michael Buszczak (68)). The obtained qPCR values (V) were normalized to those of the DNA Input and the region of cluster 1 (Cl1)/42AB as a control genomic region using the following formula: V(target)IP * V(Cl1)Input / V(Cl1)IP * V(target)Input. For each value, 3 to 7 biological replicates were analyzed. PCR primers are shown in Supplementary Table S1.

### qPCR and RT-qPCR

RNA isolation and reverse transcription was performed as described (61). Genomic DNA was isolated from the whole flies according to the standard procedure (73). Both genomic DNA and cDNAs were analyzed on DT96 real-time DNA amplifier (DNA-Technology) using SYTO™ 13 Green Fluorescent Nucleic Acid Stain (Thermo Fisher Scientific). RT-qPCR values were normalized to the rp49 mRNA and genomic DNA qPCR values were normalized to rp49 genomic qPCR. Then a ratio of RT-qPCR values to the genomic DNA qPCR values obtained with the same primers was calculated. At least three biological replicates of RNA and genomic DNA were used. All PCR-products were verified by gel electrophoresis and in some cases by sequencing. PCR primers are shown in Supplementary Table S1.

### Bioinformatic analysis

The following ChIP-seq data and corresponding Input DNA samples were analyzed: H3K9me3 and HP1a ChIP-seq from *piwi*^*2*^/+;*piwi*^*Nt*^/+ ovaries Gene Expression Omnibus (GEO) #GSE56347 (70) (designated as line 1); H3K9me3 ChIP-seq from shWhite ovaries #GSE43829 (74) (designated as line 2); H3K9me3, HP1a and H1 ChIP-seq from Control-KD OSC #GSE81434 (75). A quality control of reads and trimming of adapters for both ChIP-seq and PRO-seq data were carried out using FastQC version 0.11.5 and Trimmomatic version 0.39 (http://usadellab.org/cms/?page=trimmomatic/), respectively. Read mapping to the *D. melanogaster* reference genome (release 5.57 from flybase.org) was performed using Bowtie2 version 4.1 (http://bowtie-bio.sourceforge.net/index.shtml/) with the mapping option -k1. Further filtering and indexing were carried out using Samtools version 1.7 (http://www.htslib.org/). To create coverage profiles, the data was converted to reads per million (RPM) and the region of interest was divided into 100 nt windows. For further normalization of coverage profiles to Input DNA-seq in each window, we used bedtools version 2.26.0 (https://bedtools.readthedocs.io/en/latest/). PRO-seq data for *w*^*1118*^ (wildtype) ovaries (76) was taken from GEO #GSM3608097 and normalized to Input sample from *w*^*1118*^ ovaries #GSM3608100. To obtain transcript per million (TPM), we used Salmon version 1.3.0 (https://salmon.readthedocs.io/en/latest/salmon.html/) (77) with the size of the kmer = 31. The rDNA, R1 and R2 TPMs were additionally normalized to the numbers of copies in the genome, which were calculated by dividing corresponding RPKM (Reads Per Kilobase of transcript, per Million mapped reads) values by RPKM values of single-copy genes. The obtained TPMs of rDNA, R1, R2 and protein-coding genes were converted to log2 and visualized using the ggplot2 package (https://ggplot2.tidyverse.org/).

### Cell line and Actinomycin D treatment

Ovarian somatic cells (OSC) were grown at 25°C in Shields and Sang M3 insect medium (Sigma-Aldrich **#**S3652) supplemented with 10% heat inactivated fetal bovine serum - FBS (Gibco #10270106), 10% fly extract, 10 µg/mL insulin (Sigma-Aldrich # I9278), 0.6 mg/mL L-glutathione (Sigma-Aldrich #G6013), 50 units/mL penicillin and 50 µg/mL streptomycin. To obtain the fly extract, 1 g of flies, 3–7 days after eclosion, were homogenized in 6.8 mL of M3 medium and centrifuged for 15 min at 1500 x g. The supernatant was heated for 10 min at 60°C and centrifuged, after which the pellet was discarded. Actinomycin D (Act D) (Sigma #A9415) was used in the final concentration of 10 μg/mL in OSC growth medium for 1 h at 25°C.

### RNA FISH, immunostaining, and detection of nascent transcripts

RNA FISH with an R2 probe was performed using tyramide signal amplification as previously described (78). The DNA template for R2 probe transcription carrying the T7 RNA polymerase promoter sequence at their 5′-end was PCR-amplified on the *Drosophila* genomic DNA using primers indicated in Supplemental Table S1. Single molecule RNA FISH (smFISH) for pre-rRNA was carried out using CY5-labeling oligonucleotide probes (Supplemental Table S1). Dissected ovaries were fixed in 4% PFA, then washed in PBS (supplemented with 0.1% Tween20 and 0.3% Triton X100). Ovaries were then incubated in buffer F containing 50% formamide, 2x SSC, and 0.1% Tween20. Ovaries were briefly heated (80°C) in 100 µL of hybridization buffer containing 50% formamide, 2x SSC, 0.1% Tween20, 10 µg herring sperm DNA, 10 µg yeast tRNA, 0.1 mM DTT and 15 ng DNA labeled probes and then were incubated overnight at 37°C. Ovaries were washed in buffer F with decreasing concentration of formamide (50%, 25%, 10%) and then with PBS.

Immunostaining was performed according to previously described protocols for OSC cells (79) and ovaries (80). The following primary antibodies were used: rabbit polyclonal anti-fibrillarin (1:500, Abcam #5821); mouse monoclonal anti-fibrillarin (1:500, Abcam #4566); rabbit anti-lamin (1:500, provided by P. Fisher) (81), guinea pig polyclonal anti-Udd (1:800, provided by M. Buszczak) (68). Secondary antibodies (Invitrogen, Thermo Fisher Scientific) were the following: anti-rabbit IgG Alexa Fluor 488; anti-rabbit IgG Alexa Fluor 546; anti-rabbit IgG Alexa Fluor 633; anti-mouse IgG Alexa Fluor 488; anti-mouse IgG Alexa Fluor 546; anti-mouse IgG Alexa Fluor 633 and anti-guinea pig IgG Alexa Fluor 633.

The newly synthesized RNA in OSC cells was labeled with 5-ethynyl-uridine (EU) (Life Technologies, #E10345) as described (61). For EU incorporation in ovaries, manually dissected ovaries were incubated in the OSC growth medium with 1 mM EU for 1 h at 25°C. Then ovaries were fixed and permeabilized as described (79). EU detection using the Click-iT™ reaction was carried out in a cocktail containing Alexa Fluor 647 azide, triethylammonium salt (#A10277, Invitrogen, Thermo Fisher Scientific) and Reaction Buffer Kit (#C10269, Invitrogen, Thermo Fisher Scientific) for 30 min in the dark at room temperature. Then ovaries were washed in PBTX and processed for immunostaining. Confocal microscopy was performed using an LSM 510 META system (Zeiss).

## Results

### R1 and R2 elements are enriched in repressive chromatin marks compared to uninserted rRNA genes

First, to clarify the extent to which the inserted rRNA genes are repressed in ovaries, we evaluated levels of pre-rRNA, as well as R1 and R2 nascent transcripts in publicly available PRO-seq data for ovaries of the *w*^*1118*^ *D. melanogaster* line (76). To account for the copy number of these repetitive elements in the genome, we normalized PRO-seq values to the number of corresponding sequences in the genomic DNA sample of the same *Drosophila* strain (76). During this analysis we calculated that the *w*^*1118*^ female genome contains a total of 206 rDNA genes, of which 20 (∼10%) have R2 insertions. It should be also noted that R1 elements can insert outside the rDNA locus and therefore the R2 sequence is more appropriate as a marker of inserted rDNA. As a result, our analysis showed that the average pre-rRNA transcription level per rDNA repeat is higher than transcription of any protein-coding gene, and approximately 30 and 100 times higher than that of R2 or R1 elements, respectively (Figure 1B). In fact, transcription rates of inserted and vigorously transcribed rRNA genes can differ much stronger, given that active rRNA genes may constitute only a fraction of all uninserted rDNA units.

Next, we asked whether inserted rDNA repeats are enriched in repressive chromatin marks. In order to check this, we first re-analyzed our own (70) and publicly available ChIP-seq data from ovaries (74) and ovarian somatic cultured cells (OSC) (75) normalized to the DNA Input samples of corresponding genomes. We observed about a 2-fold enrichment of the H3K9me3 canonical heterochromatin mark, as well as a main reader of this mark, HP1a (82), in R2 and R1-associated chromatin compared to the rDNA sequence in all analyzed ChIP-seq libraries (Figure 1C). In addition, in OSC cells R1/R2 elements exhibited an increased level of H1 linker histone (Figure 1C), which also serves as repressive chromatin mark in *Drosophila* (62,75,83).

We then performed ChIP-qPCR to evaluate H3K9me3 and HP1a levels in the chromatin of inserted and uninserted rDNA units separately taking advantage of PCR to detect uninserted rRNA genes using primers surrounding the R1/R2 insertion sites in the 28S rDNA sequence (primers location is shown on Figure 1A). We found a more than 3-fold higher H3K9me3 enrichment in the chromatin of R2 insertions compared to uninserted rDNA units in the ovaries of two tested *Drosophila* lines and about a 2.5-fold higher enrichment of HP1a (Figure 1D). Thus, our results unequivocally show that inserted rDNA repeats are much more strongly associated with heterochromatin marks than rDNA units lacking insertions. Interestingly, we also observed reduced H3K9me3 and HP1a occupancies in the very beginning of ETS (Figure 1D, primer location on Figure 1A), which corresponds to promoter regions of all rDNA units (both inserted and uninserted). This result indicates that chromatin marks are non-uniformly distributed along a single rDNA repeat, and promoter regions of rDNA genes are depleted in heterochromatin marks.

### Disruption of heterochromatin only partially weakens the repression of inserted rRNA genes

To estimate the impact of repressive chromatin state on the silencing of rDNA units with TE insertions, we analyzed the levels of R1 and R2 transcripts in ovaries depleted for H3K9-specific methyltransferases and HP1a. According to RT-qPCR, nos-GAL4 driven germline knockdown (GKD) of H3K9 methyltransferase Eggless/SetDB1 led to about a 2-fold upregulation of R2 elements as compared to the control GKD of white (Figure 2A). Note that Eggless was previously shown to be strongly required for transcriptional silencing of a broad range of TEs in ovaries (84) and its knockdown caused about a 70-fold derepression of telomeric HeT-A element in our analysis (Figure 2A). GKD of another H3K9 methyltransferase, Su(var)3-9, which along with TE repression is involved in the formation of constitutive heterochromatin and the spreading of the H3K9me3 mark (84,85) induced a 2-3-fold upregulation of R2 and R1 elements (Figure 2B). In contrast to previous report (86), we found that Su(var)3-9 GKD females were sterile and exhibited much more severe defects of oogenesis than Eggless-depleted individuals. Similarly, the level of R2 transcripts increased approximately 3-fold upon HP1a GKD (Figure 2C), which also induced a defective ovarian phenotype, although HP1a-deficient germ cells survived the first days after birth according to immunostaining.

**Figure 2.**
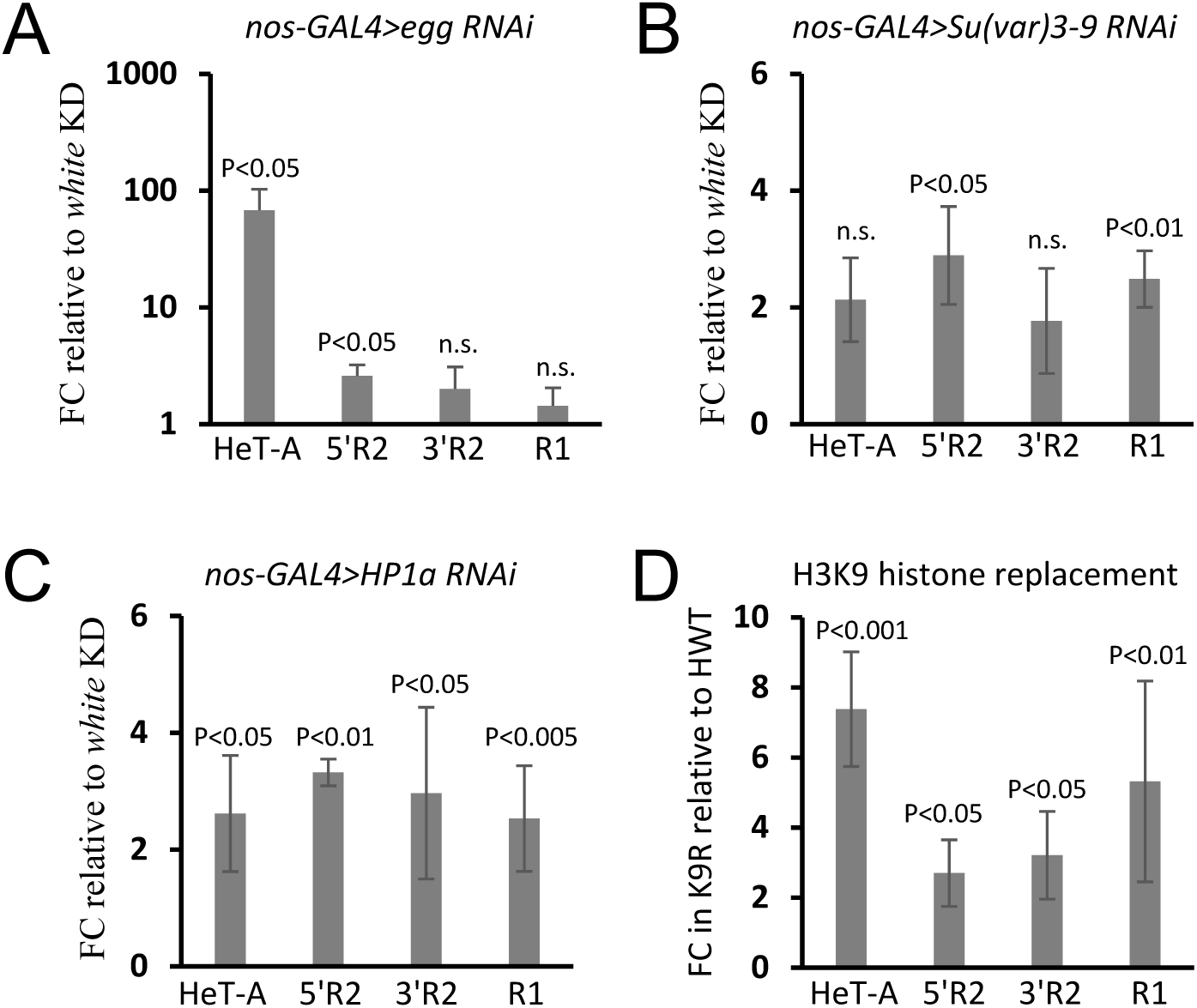
Lack of heterochromatin components exerts a moderate effect on expression of inserted rRNA genes. **(A-C)** RT-qPCR analysis of HeT-A, R1 and two regions (5’ and 3’) of R2 elements in ovaries upon nos-GAL4 driven germline knockdown (GKD) of Eggless/SetDB1 (**A**), Su(var)3-9 (**B**) and HP1a (**C**) proteins. Mean fold change (FC) +/−s.d. relative to the white GKD normalized on the rp49 transcript levels are indicated. (**D)** RT-qPCR analysis of HeT-A, R1 and R2 elements in larvae lacking H3K9 methylation. FC +/−s.d. in larvae with replacement of H3K9 lysine residues with arginine (K9R) relative to control individuals containing a transgenic wild-type histone array (HWT) and normalized to rp49 is shown. p-values are based on two-sided Student’s t-test.

Then, to directly evaluate the role of H3K9 methylation in the repression of inserted rDNA units we took advantage of a recently developed histone replacement system (71). In this system, a wild-type cluster of histones can be replaced by an artificial one, in which codons for K9 lysine residues of the H3 histone genes are substituted with arginine (K9R) (see Materials and Methods for details). Despite the fact that K9R individuals are almost devoid of the H3K9me2/3 heterochromatin histone modifications they survive up to the larval stage (71). RT-qPCR analysis of female larvae showed that K9R substitution induced an approximately 3-fold increase of R2 expression as compared to control individuals (HWT) containing a wild-type histone transgenic array, which rescues the histone deletion (Figure 2D). Some upregulation of R1 elements was also observed in K9R larvae (Figure 2D). Thus, the loss of heterochromatin components causes a 2-3-fold increase of R1/R2 transcription that is much less than the derepression that would be expected if the silencing of inserted rRNA genes was disrupted. These results suggest that the putative mechanisms of individual rDNA unit repression mostly retain their effectiveness in the absence of heterochromatin marks.

### Udd is required for silencing of R2-inserted rDNA units in germ cells

It is possible that the process of selective silencing/activation of individual rDNA units is coupled with the Pol I transcription initiation apparatus. M. Buszczak with coworkers described the *Drosophila* Selectivity Factor I like (SL1-like) protein complex that is essential for the transcription initiation by Pol I on rDNA promoters (68). Along with the conserved proteins TAF1B and TAF1C-like, which have homologues in the mammalian SL1 complex, another *Drosophila*-specific component of this complex called Under-developed (Udd) was identified. The authors showed that the loss of Udd compromised rRNA synthesis and development of ovarian germ cells and caused sterility (68).

We checked the effects of SL1-like components on R2 expression by RT-qPCR and unexpectedly revealed a drastic (70-80-fold) upregulation of R2 elements in *udd*^*1*^*/udd*^*1*^ ovaries, as well as in *udd*^*1*^*/udd*^*null*^ ovaries, relative to corresponding heterozygotes (Figure 3A). Although Udd is expressed ubiquitously and was detected within nucleoli of various *Drosophila* tissues (Supplementary Figure S1), we did not observe any significant influence of *udd* mutations on R2 in carcasses (bodies without gonads), while about a10-fold derepression of R2 elements was found in testes of *udd* mutant males (Figure 3A). R1 elements were also significantly upregulated in *udd*^*1*^*/udd*^*1*^ and *udd*^*1*^*/udd*^*null*^ ovaries, although weaker than R2. Expression of other analyzed TEs showed a smaller increase (HeT-A) or no change (Zam) (Figure 3A). Next, we examined in which types of ovarian cells R2 activation occurs in *udd* mutants. *Drosophila* ovaries are composed of ovarioles, chains of egg chambers starting from a germarium region and then consistently developing over 14 stages. Each egg chamber includes 16 cytoplasmically connected germline cells (fifteen nurse cells and a single oocyte) surrounded by somatic follicle cells. Nurse cells are polyploid (up to 8000C) and exhibit high transcription activity supplying the transcriptionally inert oocyte with proteins, RNA and ribosomes through intercellular channels (87). RNA-FISH using a probe targeting the 3’-region of the R2 element combined with immunostaining for fibrillarin nucleolar marker revealed an accumulation of R2 transcripts in the nucleoli of nurse cells, but not in somatic cells of *udd*^*1*^*/udd*^*null*^ ovaries (Figure 3B). In the germ cells of germaria and stage 2 egg chambers only a weak FISH signal was observed, which then increased as oogenesis progressed and reached a maximum in nurse cells at stages 4-5, whereas the later stages were absent in *udd*^*1*^*/udd*^*null*^ ovaries. In the control heterozygotes, the FISH signal was not detected in any cells (Figure 3B). It is noteworthy that in nurse cells nucleoli develop into branched structures different from the ordinary ring-shaped nucleolus (67), but this does not occur in *udd*^*1*^*/udd*^*null*^ ovaries (Figure 3B).

**Figure 3.**
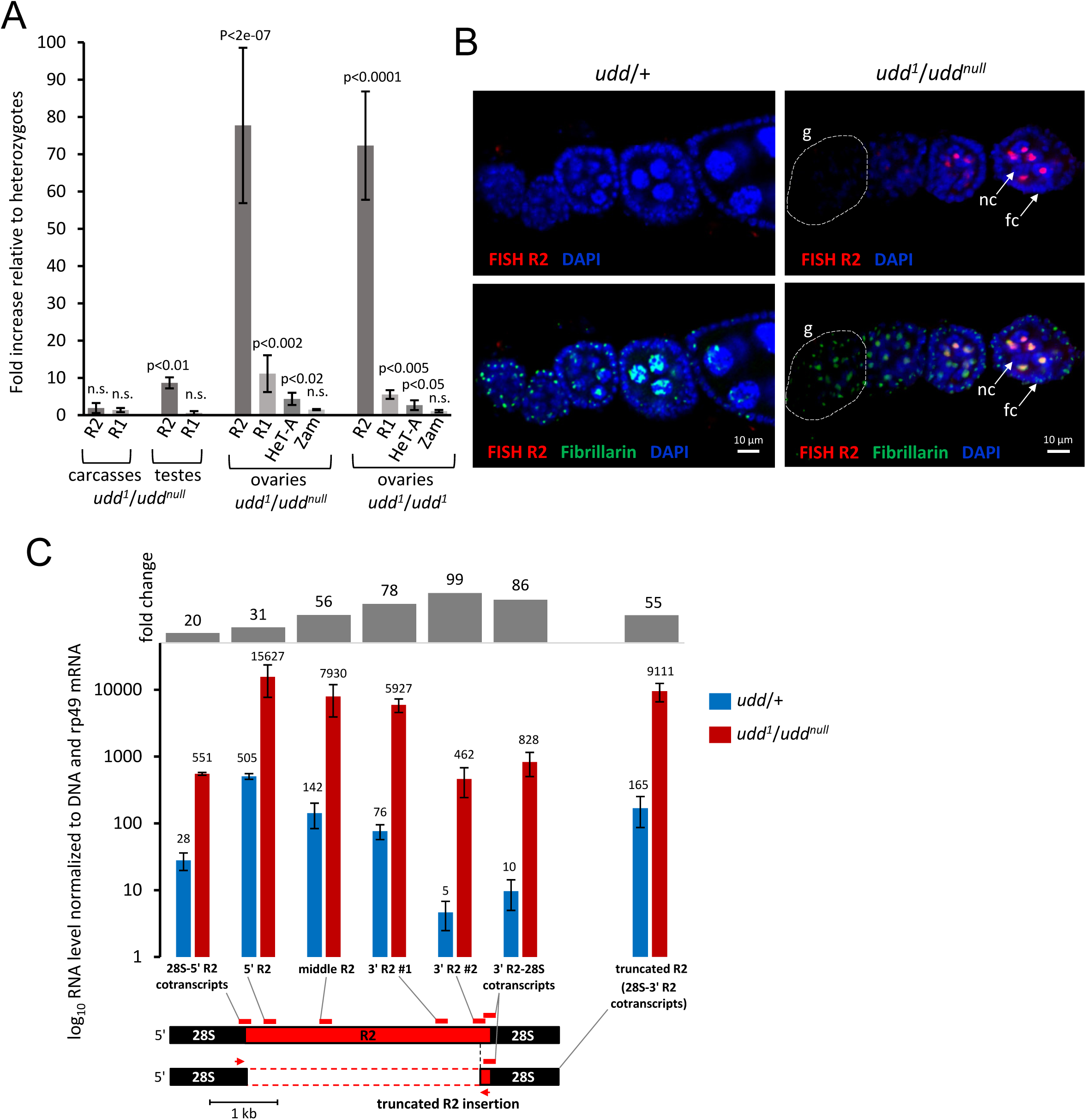
Udd is required for silencing of R2-inserted rDNA units in germ cells. **(A)** RT-qPCR analysis of R1 and R2 elements in ovaries, testes, and carcasses of *udd* mutants. Mean fold increase +/−s.d. relative to the corresponding heterozygous sisters normalized on the rp49 transcript levels are indicated. p-values from Student’s t-test for differences between homo(trans-hetero)zygotes and heterozygotes are shown. n.s. = not significant. (**B)** RNA-FISH of R2 transcripts (red) combined with immunostaining for nucleolar marker fibrillarin (green) in ovarioles of *udd*^*1*^/*udd*^*null*^ mutants and control heterozygotes (*udd*/+). A germarium regions are indicated as “g” and circled with a dotted line; examples of germline nurse cells and somatic follicle cells are indicated as “nc” and “fc”, respectively. (**C)** Fold change and log10 RNA level normalized to genomic DNA and rp49 mRNA measured by RT-qPCR in *udd*^*1*/^*udd*^*null*^ (red bars) and control *udd*/+ (blue bars) ovaries. Location of analyzed PCR fragments on the sequence of the 28S gene containing the R2 insertion is indicated by red rectangles. Location of primers used for detection of transcripts corresponding to rRNA genes with truncated R2 insertions is shown by red arrows. Note that 3’ R2-28S cotranscripts correspond to both full-length and truncated R2 elements. Since 3’-ends of the truncated R2 inserts are transcribed more efficiently than in full-length elements, the level of 3’ R2-28S cotranscripts is higher than that of 3’ R2 #2 region, which does not correspond to the truncated R2 copies.

Our results challenged the previous finding that Udd serves as a component of SL1-like complex necessary for rDNA transcription initiation (68) and implied it’s another function in discriminating between inserted and uninserted rDNA copies in germ cells. Udd is small (18 kD) protein and does not contain any known motifs. Although components of the Pol I transcription apparatus are usually highly conservative (1), Udd has no homologs outside the *Diptera*. Moreover, *udd*^*1*^*/udd*^*1*^ and *udd*^*1*^*/udd*^*null*^ flies do not display any morphological features of reduced ribosome production (minute-like phenotypes) (88), whereas they are sterile and have severely reduced ovaries and testes. These facts prompted us to re-check the effect of Udd on total rRNA synthesis in ovaries. Detection of pre-rRNA transcripts using smFISH demonstrated the presence of rDNA transcription in all cell types and in all oogenesis stages, which are retained in *udd*^*1*^/*udd*^*null*^ mutants (Supplementary Figure S2A). Similar results were obtained using detection of nascent RNA by EU-incorporation analysis (Supplementary Figure S2B). Nevertheless, it cannot be ruled out that cells with a strong decline in rRNA level die, which leads to the observed elimination of germ cells at later developmental stages and their loss with age. Quantification of pre-rRNA levels by RT-qPCR showed about a 2-fold decrease of 18S-ITS1 cotranscripts in *udd*^*1*^/*udd*^*null*^ ovaries relative to heterozygotes (Supplementary Figure S2C). Thus, in the absence of Udd, total rRNA synthesis in the germline can be reduced, albeit to a lesser extent than previously shown (68), whereas there is a strong overexpression of normally silent rDNA units.

Previously it was suggested that repression of rDNA repeats with TE insertions can occur due to termination of transcription within the insertion sequence (50), or due to the transcriptional silencing of the entire rDNA unit (51). To test which of these repression modes is affected by Udd, we measured the amounts of transcripts corresponding to different regions of R2 insertions in the *udd/+* and *udd*^*1*^/*udd*^*null*^ ovaries. Then we normalized the RT-qPCR values to *udd* genomic DNA qPCR values obtained using the same primers. Therefore, the results shown in Figure 3C allow us to compare the transcript levels between different regions of R2 regardless of their abundance in the genome and regardless of primer amplification efficiency. The normalized RT-qPCR values were drastically higher in the *udd*^*1*^/*udd*^*null*^ ovaries than in the control for all analyzed regions indicating increased transcription along the entire length of R2 insertions (Figure 3C). Importantly, we observed an approximately 20-fold upregulation of transcripts containing the junction of the upstream 28S rRNA and the beginning of the R2 sequence (hereafter, “28S-5’R2 cotranscripts”) (Figure 3C). Thus, derepression occurs at least partially due to the enhancement of rDNA transcription upstream of the insertion region. Note that the absolute level of 28S-5’R2 cotranscripts is lower than that of 5’ R2 body transcripts (Figure 3C), because R2 RNA can self-cleave from the 28S rRNA sequence. At the same time, we observed a reduction of the transcript amount in the 5’-to 3’ direction of the R2 body which was especially prominent at its very 3’-end (3’ R2 #2 region on Figure 3C). In *udd*/+ ovaries, the amount of transcripts detected in the 5’-region of the R2 was approximately 100-fold higher than in the 3’-region (Figure 3C). In *udd*^*1*^/*udd*^*null*^ ovaries, this difference was only slightly smoothed out and amounted to about 30 times (Figure 3C). Altogether, these results suggest that normally repression of R2-inserted rRNA genes occurs both due to silencing of the entire rDNA unit, as well as by interruption of transcription in the body of R2, whereas Udd is basically required for the repression of the entire rRNA gene. The loss of Udd strongly enhances the passage of transcription across rDNA unit with R2 insertion and additionally slightly facilitates transcription within the R2 body.

Along with full-length R2 elements, some rDNA units can contain R2 insertions shortened to varying degrees from the 5’-end, formed due to abortive reverse transcription (30,41). Using genomic DNA PCR with subsequent sequencing, we detected the presence of highly truncated R2 inserts of ∼180 and ∼50 bp in the *udd*^*1*^/*udd*^*null*^ genome, whereas full-length R2 is about 3.6 kb. RT-qPCR using a forward primer located in upstream 28S rRNA sequence and reverse primer in the 3’-end of R2 demonstrated about a 50-fold upregulation of rDNA units with these truncated R2 inserts in *udd*^*1*^/*udd*^*null*^ ovaries that is similar to the derepression of those with full-length R2 elements (Figure 3C). Thus, Udd’s effect on inserted rDNA units is independent of the insertion length and therefore is unlikely to be determined by any specific nucleotide motifs within the R2 sequence, or at least within its most part. Moreover, this result further supports the conclusion that loss of Udd causes transcriptional activation of the entire rDNA unit containing insertion.

### Udd is associated with transcriptionally active, but not silent rDNA repeats

Based on our results, it is formally possible that Udd can serve as a component of a hypothetical repressor complex interacted with silent rDNA units. Consistent with a previous report (68), our ChIP-qPCR analysis of *Batumi* wild type ovaries showed high Udd enrichment at the rDNA promoter regions (ETS on Figure 4A), which correspond to both inserted and uninserted rDNA repeats. However, we also observed a weak but statistically significant association of Udd with uninserted 28S rRNA sequences that was about 2-fold higher compared to different regions of R2 insertions (Figure 4A). The same Udd enrichment at uninserted 28S as compared to R2 was detected in ovaries of two other *D. melanogaster* lines (Supplementary Figure S3). This effect can be attributable to the preferential interaction of Udd with actively transcribed rRNA genes. Studies on mammalian cells show that rDNA repeats in the nucleolus form loops connecting promoter and termination regions (89,90) or more complex solenoid-like structures, in which the transcribed rDNA region is cylindrically wrapped around the core formed by SL1 (91). Thus, it is conceivable that in ChIP experiments some cross-linking can occur between proteins associated with the promoter and pre-rRNA-coding part of the same rDNA unit. Alternatively, Udd can spread along the vigorously transcribed rRNA gene. We also found Udd enrichment in the region of rDNA transcription termination on the border of 28S and IGS sequences, as well as in the IGS (Figure 4A; Supplementary Figure S3). This can be interpreted as the formation of a spatial loop or as a binding of the SL1-like complex to putative IGS promoters. A similar pattern of Udd distribution along rDNA units was observed in both ovaries and carcasses (Supplementary Figure S3). However, enrichment levels were lower in carcasses, which may reflect a higher rate of rRNA synthesis in ovarian cells.

**Figure 4.**
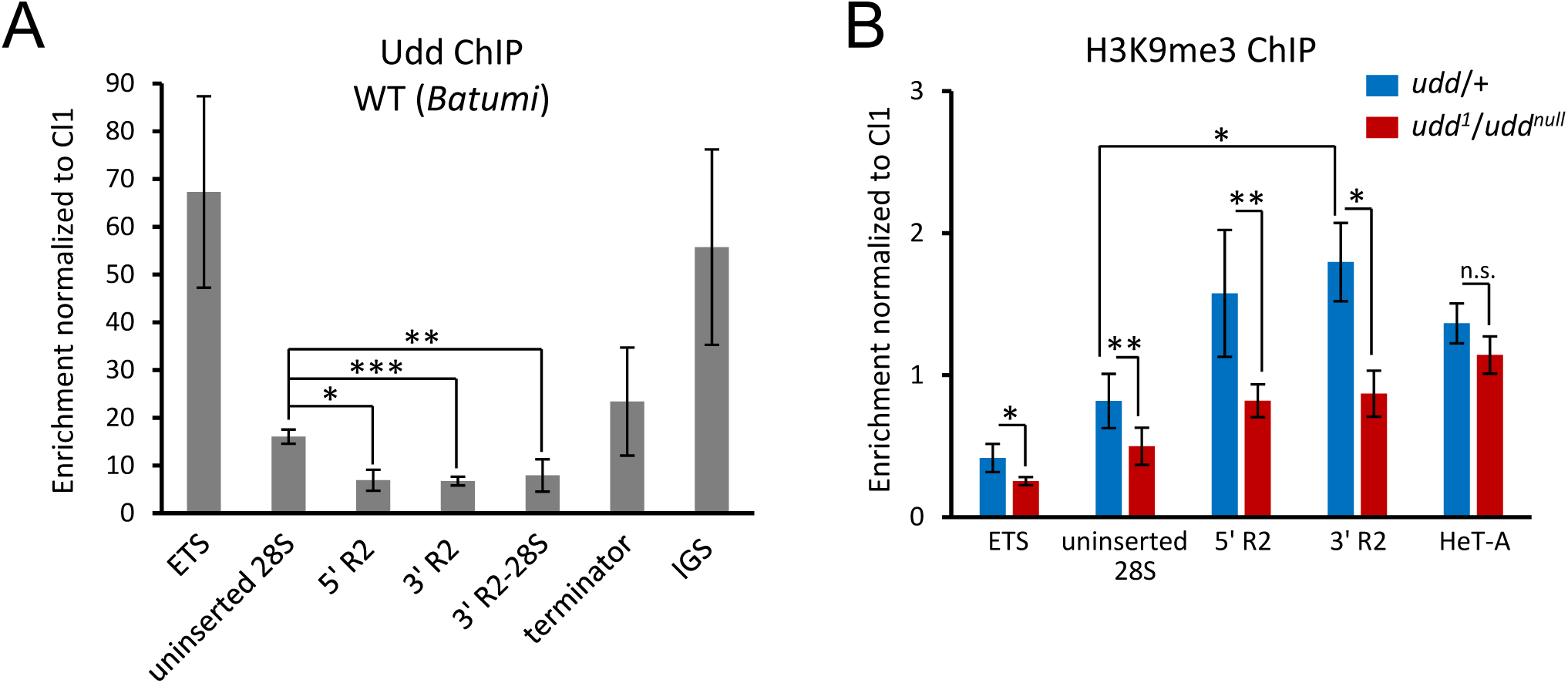
Udd indirectly controls chromatin status of R2 insertions. **(A)** ChIP-qPCR analysis of *Batumi* wild type ovaries showing Udd enrichment levels at different regions of rDNA repeats: beginning of ETS; uninserted 28S sequence; 5’ and 3’ regions of R2 insertions; the border of R2 3’-end and 28S sequences (3’ R2-28S); the border of 28S and IGS sequences (terminator); 330bp IGS repeat (IGS). (**B)** ChIP-qPCR analysis of H3K9me3 level in *udd*^*1*/^*udd*^*null*^ (red bars) and control *udd*/+ (blue bars) ovaries. Mean +/− s.d. and p-values based on Student’s t-test are indicated, * p<0.003, ** p<0.01, *** p<0.0005, n.s. = not significant.

### Udd affects chromatin status of rDNA repeats

We tested whether derepression of R2-inserted rDNA units upon the Udd loss was accompanied by changes in repressive chromatin marks. ChIP-qPCR revealed 2-fold reduction of H3K9me3 occupancy in the chromatin of R2 elements and rDNA promoter regions in *udd*^*1*^/*udd*^*null*^ ovaries when compared to control heterozygotes (Figure 4B). Unexpectedly, the H3K9me3 level in the chromatin associated with uninserted 28S was also noticeably reduced, though to a lesser extent when compared to the R2 sequence (Figure 4B). Based on this result, it is tempting to speculate that some fraction of uninserted rDNA genes can be also repressed normally, whereas upon the Udd loss they are transcriptionally activated and lose heterochromatin marks in a similar way to inserted rRNA genes.

We found no effect of *udd* mutation on the H3K9me3 occupancy in the chromatin associated with telomeric *HeT-A* elements (Figure 4B) and other heterochromatic genome regions (data not shown), indicating that Udd controls the H3K9me3 recruitment specifically within rDNA locus. We also observed a reduction of HP1a in the chromatin associated with both R2 insertions and uninserted 28S rDNA sequences in *udd*^*1*^/*udd*^*null*^ ovaries (Supplementary Figure S4). However, the levels of H3K27me3 and H4K20me3 marks, which have been shown to be involved in rDNA silencing in mammalian cells, along with H3K9me2/3 marks (92,93) did not change significantly for any of the analyzed sequences (Supplementary Figure S4).

### Inhibition of transcription erases epigenetic differences between inserted and uninserted rDNA units

Our observations suggest that the reduction of H3K9me3 occupancy observed in rRNA genes in *udd* mutant ovaries may be due to their transcriptional activation, and vice versa normally repressive chromatin marks may be recruited to rDNA units as a consequence of their silencing. To test this assumption, we examined changes in rDNA chromatin after inhibition of transcription by actinomycin D (ActD) in OSC cultured cells. ActD nonspecifically blocks activity of all types of RNA polymerase (94). Cessation of rRNA synthesis in ActD-treated cells was detected by the absence of EU incorporation into nascent transcripts in the nucleolus, which was accompanied by a reduction of a nucleolar area occupied by fibrillarin (Figure 5A). Interestingly, Udd was largely released from the nucleolus and found in the cytoplasm of ActD-treated cells (Figure 5A). Consistent with this, ChIP-qPCR revealed a loss of Udd binding in the rDNA promoter, terminator, and IGS regions where Udd is enriched in untreated cells (Figure 5B).

**Figure 5.**
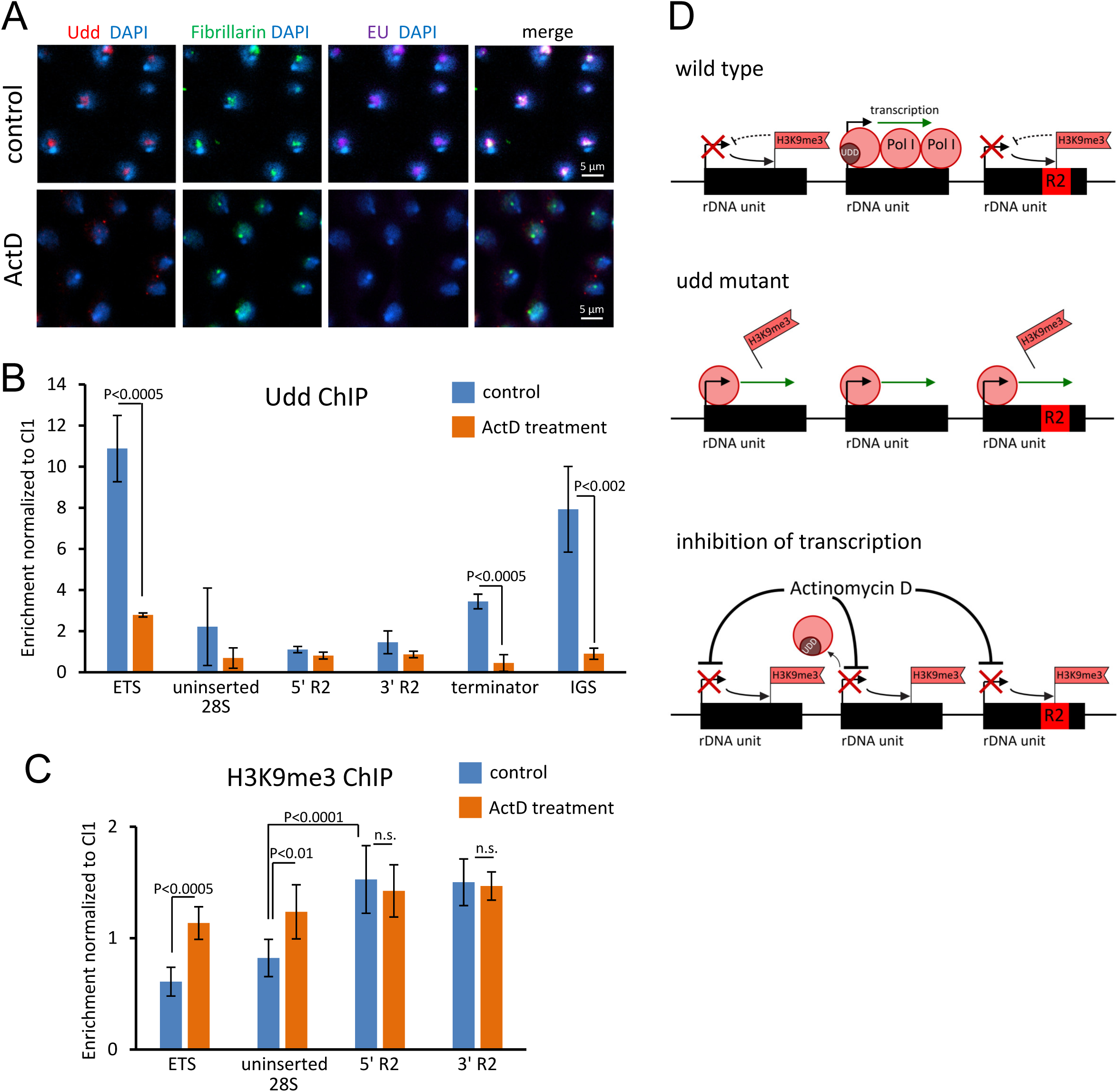
Inhibition of transcription leads to recruitment of H3K9me3 to uninserted rDNA repeats. **(A)** Control and actinomycin D (ActD)-treated OSC cells immunostained for Udd, fibrillarin and counterstained for nascent RNA (5-Ethynyl-uridine (EU) incorporation) and DAPI. (**B)** ChIP-qPCR analysis of Udd level in control and ActD-treated OSC cells at the beginning of ETS, uninserted 28S sequence, R2 insertions, the border of 28S and IGS sequences (terminator) and 330bp IGS repeats (IGS). Mean +/−s.d. and p-values based on Student’s t-test are indicated. (**C)** ChIP-qPCR analysis of H3K9me3 mark in control and ActD-treated OSC cells. p-values are calculated using Student’s t-test, n.s. = not significant. (**D)** A working model for the role of Udd and heterochromatin components in selective silencing of rDNA repeats. Upper panel: in wild-type ovarian cells, Udd is required for accumulation of the Pol I transcription complexes at some uninserted rDNA units possibly due to facilitating transcription re-initiation. rDNA genes with insertions and the rest of uninserted rRNA genes remain less transcriptionally active that leads to establishment of H3K9me3 mark. The heterochromatic state further enhances repression. Middle panel: the loss of Udd leads to redistribution of Pol I complexes from active uninserted rRNA genes to all rRNA genes. Transcriptional activation of normally silent rDNA units causes the reduction of H3K9me3 level in their chromatin. Bottom panel: upon actinomycin D treatment, transcription of all genes is stopped. This leads to the removal of Udd from promoters and uniform establishment of H3K9me3 on all rDNA units.

Examination of H3K9me3 levels showed that, as in the ovaries, inserted rDNA genes were noticeably enriched in H3K9me3 compared to both uninserted ones and promoter regions in control cells (Figure 5C). ActD treatment induced a significant increase of H3K9me3 levels in the chromatin corresponding to uninserted 28S rDNA sequence, as well as to the rDNA promoter region, but not to R2 insertions (Figure 5C). This led to the disappearance of the differences between H3K9me3 occupancies in the chromatin of the analyzed DNA fragments (Figure 5C). Thus, in the absence of transcription, uninserted rDNA units acquire the same chromatin status as normally silent inserted ones. As a result, an approximately uniform level of heterochromatinization is established throughout the entire rDNA cluster. Although it cannot be ruled out that the observed changes can be determined by more general processes caused by the nucleolar stress, the results support the model that the differences in H3K9me3 levels between inserted and uninserted rRNA genes is a consequence of their different transcription levels.

## Discussion

### Possible mechanisms of discrimination between inserted and uninserted rDNA units

It has been known for several decades that a substantial number of rDNA repeats in the *Drosophila* genome contain insertions and that such genes are usually not transcribed or are transcribed at a very low level (47,48,51). Here we found that a mutation in a single gene, *udd*, can disrupt this silencing, inducing almost 2 orders of magnitude upregulation of R2-containing rDNA units in ovarian nurse cells (Figure 3). This effect is also accompanied by a moderate reduction of overall pre-rRNA production (Supplementary Figure S2). Thus, the loss of Udd likely impairs a mechanism normally discriminating between rRNA genes selected for active transcription and silenced repeats: active uninserted genes begin to be transcribed weaker, while silent genes with R2 insertions become drastically upregulated.

What assumptions can be made about the molecular function of Udd protein? Generally, our observations fit well with the idea that Udd is associated with the Pol I transcription initiation apparatus during active transcription of rRNA genes. First, Udd has previously been shown to interact with conservative components of SL1 complex (68). Secondly, we found a preferential Udd association with uninserted rRNA genes (Figure 4A; Supplementary Figure S3). Thirdly, inhibition of transcription resulted in the removal of Udd associated with the rDNA promoter region (Figure 5B). However, the fact that the loss of Udd does not lead to a drastic cessation of pre-rRNA synthesis (Supplementary Figure S2) suggests that Udd rather has some modulatory, but not “core” function in Pol I transcription initiation. Thus, we suggest that a choice of particular rDNA unit for transcriptional activation in ovarian cells is coupled with Pol I transcription initiation machinery. This implies that normally the initiation of transcription of the inserted rDNA units can be less efficient than those of uninserted ones. Although we could not directly test the efficiency of transcription initiation on promoters of inserted rDNA repeats, this model is indirectly confirmed by our observation that Udd loss caused an increase of R2 transcription along its entire length, including the upstream 28S rRNA sequence (Figure 3C). However, our results do not contradict a previous observation that transcription often terminates within insertions in rDNA units (50). Consistent with this work, we revealed a sharp reduction in the amount of transcripts found in the 3’-region of R2 insertions as compared to the R2 5’-region. Interestingly, this partial termination of transcription mostly retained in *udd* mutants (Figure 3C). Thus, repression of R2-inserted rDNA units in ovarian cells is likely provided by two independent means, which operate at both transcription initiation and elongation/termination levels. The first mode of silencing is Udd-dependent, whereas the interruption of transcription within R2 body is determined by other factors.

The observed Udd association with uninserted rDNA units (Figure 4A; Supplementary Figure S3) allows us to hypothesize that Udd can be involved in the redistribution of transcriptional factors and Pol I towards active rDNA repeats rather than acting as a component of a repressor complex interacting with silent units. An important question is therefore, how can the transcription initiation apparatus distinguish between inserted and uninserted rRNA genes? Mechanistically, the existence of such “transcriptional selection” means that information about the presence of R2 insertion can be transferred to the rDNA promoter, which is more than 6 kb distant from insertion. It is tempting to speculate that these interactions can be mediated by spatial loops formed between the rDNA promoter and terminator regions (89-91), which can contribute to efficient transfer from termination to re-initiation of transcription. Since transcription usually reaches the terminator in uninserted rRNA genes, it can be effectively re-initiated at their promoters. This process can potentially provide an extremely high density of Pol I complexes, as observed in active pre-rRNA-coding regions by electron microscopy (for example, (51,52). Transcription re-initiation at inserted genes should occur much less frequently due to the often termination in the R2 body. The possible role of spatial interactions in selective repression of rDNA is also supported by the observation of an 8-fold upregulation of R2 upon mutation of the chromatin architectural protein CTCF (53). It is conceivable that Udd can facilitate effective transcription re-initiation on active rDNA units or, by analogy with the mouse TTF-1 (89), it can participate in the bridging of the rDNA promoter and terminator, which is consistent with Udd enrichment at both regions (Figure 4A; Supplementary Figure S3). Within this model, the loss of Udd can release a pool of Pol I usually circulating on active rDNA genes. As a result, the released Pol I complexes are promiscuously recruited at promoters of all rDNA units, that leads to an increase in transcription of inserted rDNA genes and a weakening of uninserted ones (Figure 5D). Further experiments are needed to test these hypotheses.

The mechanisms driving inserted rDNA silencing seem to be more puzzling considering that their activation upon Udd loss occurs exclusively in germ cells (Figure 3B). This may be attributed to the fact that germ cells provide an especially high level of rRNA synthesis. It has been calculated that a stage 14 oocyte contains approximately 2 x 10^10^ ribosomes (66), the vast majority of which are synthesized in the nurse cells during stages 7 through 10 (67). Interestingly, earlier observations showed that the proportion of R2 transcripts is much higher in ovarian tissue than in embryos, larvae or pupae (49). It is plausible that Udd-based selection of rDNA units is redundant in somatic cells, where other mechanisms for the repression of inserted rRNA genes may exist. However, in nurse cells, rDNA units with insertions may overcome these silencing mechanisms due to a high abundance of Pol I transcription factors, and therefore Udd-dependent regulation becomes critical.

### rDNA transcription and heterochromatin marks

Silent rRNA genes in various organisms are known to be marked by DNA hypermethylation and regular nucleosomes carrying repressive histone marks, such as H3K9me3, H4K20me3 and H3K27me3 (92,93,95,96) (see (26-29) for reviews). Previously, heterochromatin components in *D. melanogaster* have been implicated in maintaining the nucleolar structure and preventing recombination between rDNA repeats (22). Moreover, H3K9 methylation by Su(var)3-9 is involved in nucleolar dominance, a phenomenon whereby an entire rDNA cluster is silenced (in particular, X-chromosome rDNA inactivation in *Drosophila* males) (97). However, the association of heterochromatin marks with silencing of individual rDNA units in *Drosophila* was not evident. Here we found much higher levels of H3K9me3 and HP1a in the chromatin of inserted rDNA units compared to uninserted ones (Figure 1C, 1D, 4B, 5C). Furthermore, upregulation of R2-inserted rRNA genes in the ovaries of *udd* mutants was accompanied by a reduction of H3K9me3 and HP1a enrichments (Figure 4B, S3). These results demonstrate that these marks indeed are associated with rDNA repression in *Drosophila* and provide experimental support for the idea of chromatin-based differentiation between individual rDNA repeats within one rDNA array. This allows us to draw parallels with the results obtained for human cells, where the coexistence of methylated inactive and unmethylated rRNA genes within a single rDNA cluster was demonstrated using FISH and immunostaining of single DNA fibers (20).

Another important question is whether the heterochromatin structure serves as a marker of inactive rRNA genes or it has a substantial repressive effect on their transcription. We showed that the lack of HP1a protein, H3K9 histone methyltransferases, as well as H3K9 methylation itself leads to an approximately 2-3-fold upregulation of inserted rDNA repeats in ovaries (Figure 2). These effects are quantitatively similar to the activation of R2 expression observed previously upon knockdown of histone H1, another component of heterochromatin (62). Thus, at least in ovarian cells, heterochromatinization of inserted rDNA units does not contribute as much to their repression as the Udd-dependent regulation. On the other hand, our data indicate that the level of rDNA transcription can determine its chromatin state. This is supported by the observed reduction of repressive chromatin marks at rDNA genes in *udd* mutant ovaries (Figure 4B). Furthermore, we showed that inhibition of transcription by actinomycin D treatment smooths out epigenetic differences between rRNA genes by increasing of H3K9me3 level at uninserted rDNA units (Figure 5C). These results allow us to propose that heterochromatin marks are recruited to already silent, or less effectively transcribed rDNA units, maintaining and strengthening their repression, whereas transcriptional activation can lead to removing of the repressive chromatin modifications (Figure 5D). Interestingly, in murine cells, the loss of Upstream Binding Transcription Factor (UBF), a key mammalian Pol I transcription pre-initiation factor was accompanied by recruitment of H3K9me3, histone H1 and HP1a to rDNA repeats (98). Thus, impairment of rDNA transcription initiation can lead to the establishment of repressive chromatin marks on rDNA in mammalian cells suggesting that these processes can share basic features in various organisms.

Our study opens the way to elucidating the mechanisms of transcriptional regulation of individual units within arrays of genes with identical promoters and brings us closer to understanding how their quality control occurs.

## Acknowledgments

We thank Michael Buszczak for *udd*^*1*^ and *udd*^*0*^ fly stocks and Udd antibodies, Robert J. Duronio for flies used for H3K9 histone replacement; Sarah Elgin for HP1a antibodies; Yuri Y. Shevelyov for helpful comments on the manuscript. The work was carried out with the use of the equipment of the common use center «Center of Cell and Gene Technology», Institute of Molecular Genetics, RAS.

## Figure legends

**Supplementary Figure S1.**
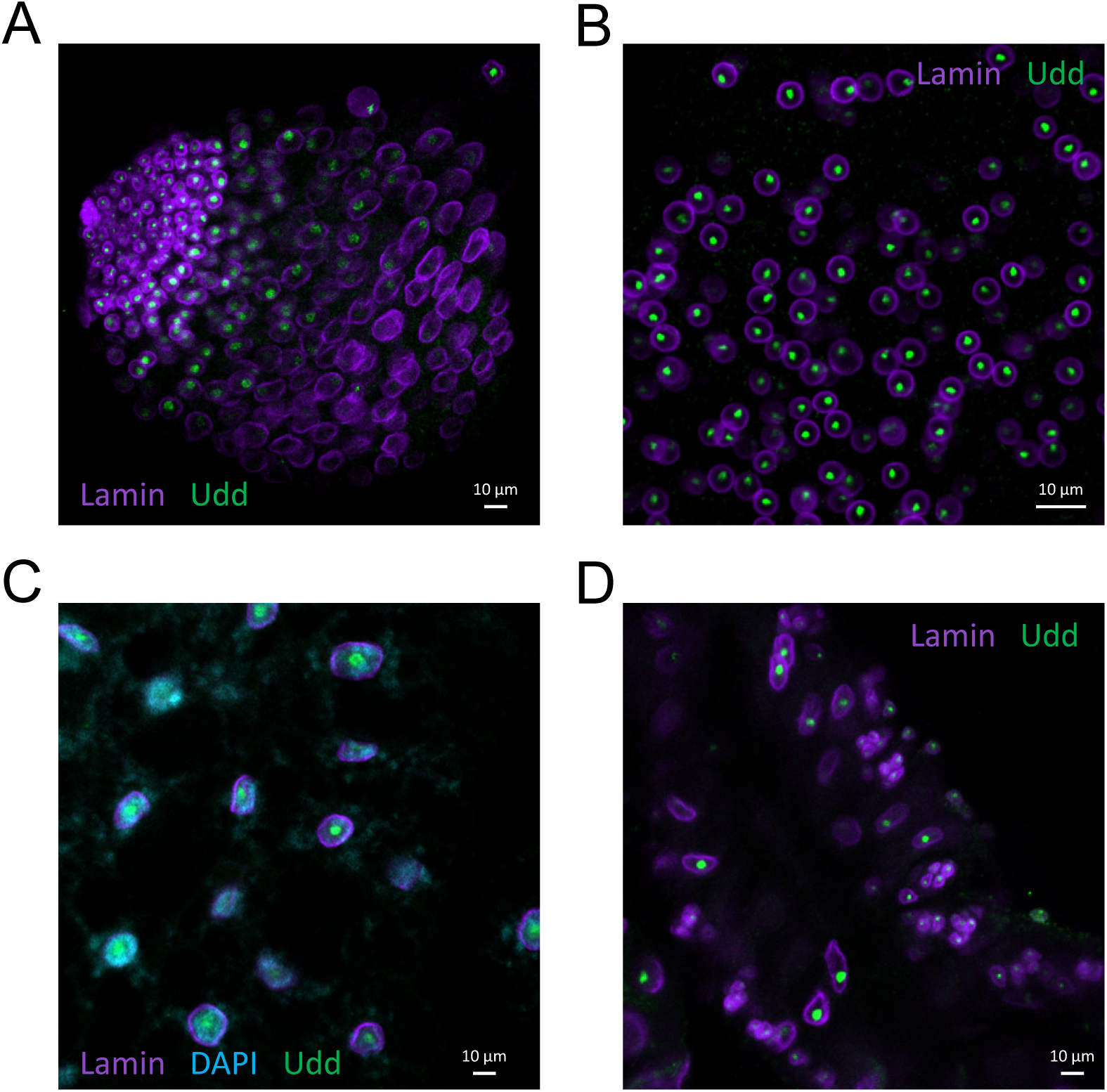
Udd localization in different tissues. Immunostaining for Udd (green) and lamin (purple) showing nuclear envelope. (**A)** Larval testes. (**B)** Adult testes accessory glands. (**C)** Fat body. (**D)** Salivary glands.

**Supplementary Figure S2.**
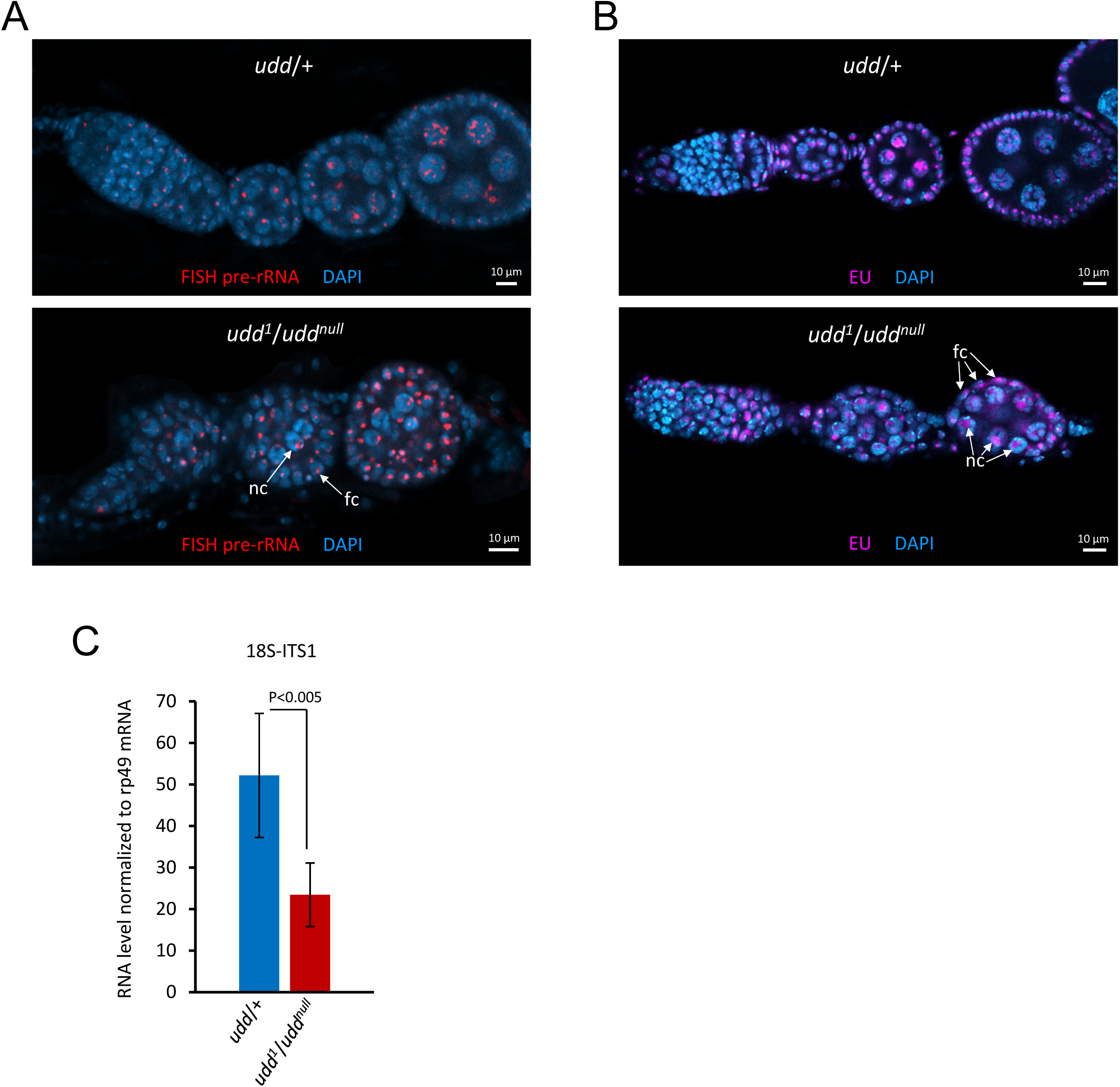
The effect of Udd loss on total rRNA synthesis in ovaries. **(A)** smFISH of pre-rRNA transcripts in *udd*^*1*^/*udd*^*null*^ and *udd*/+ ovarioles. Examples of germline nurse cells and somatic follicle cells are indicated as “nc” and “fc”, respectively. (**B)** Detection of nascent transcription by EU incorporation assay in *udd*^*1*^/*udd*^*null*^ and *udd*/+ ovarioles. Examples of EU-positive nucleoli in “nc” and “fc” are indicated by arrows. (**C)** Quantification of the pre-rRNA level (18S-ITS1) by RT-qPCR in *udd*^*1*^/*udd*^*null*^ and *udd*/+ ovaries. Mean +/−s.d. and p-value based on Student’s t-test are indicated.

**Supplementary Figure S3.**
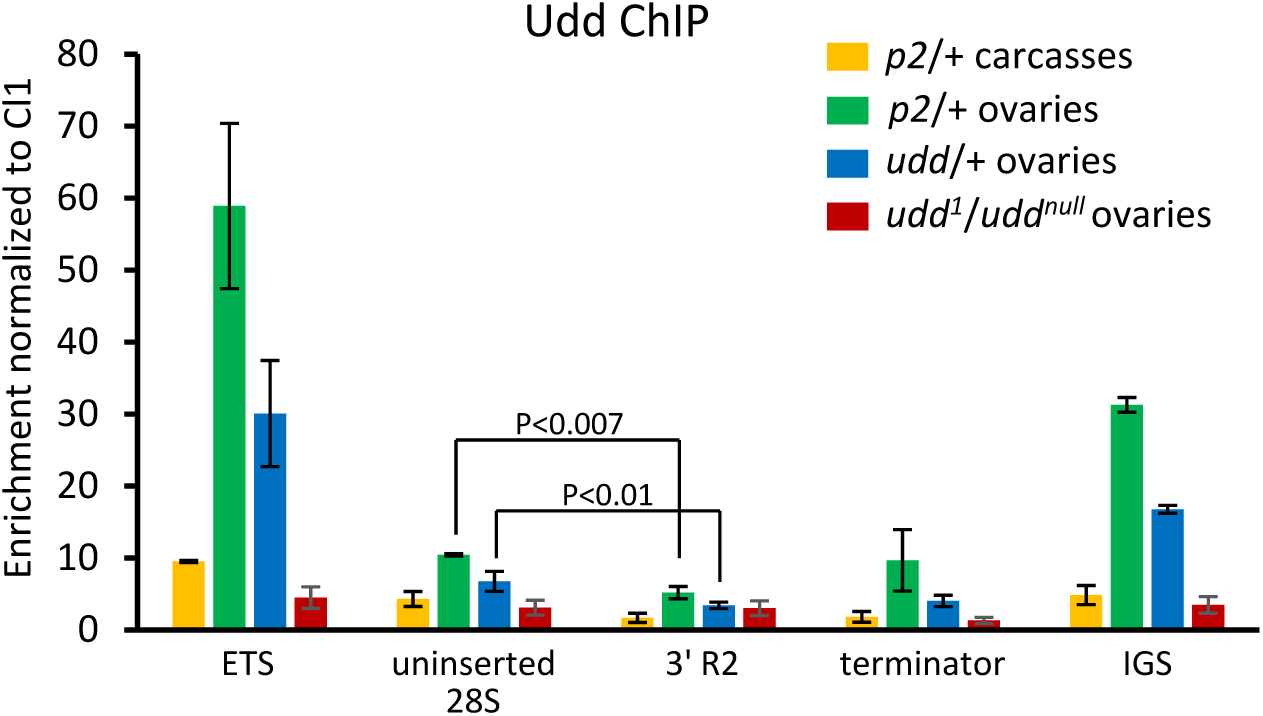
Udd ChIP-qPCR analysis of ovaries and carcasses of *piwi*^*2*^*/+* (*p2/+*) line and ovaries of *udd/+* and *udd*^*1*/^*udd*^*null*^ flies. Udd binding at different regions of rDNA repeats is shown: beginning of ETS; uninserted 28S sequence; R2 insertions; the border of 28S and IGS sequences (terminator); 330bp IGS repeat (IGS). Mean +/− s.d. and p-values based on Student’s t-test are indicated.

**Supplementary Figure S4.**
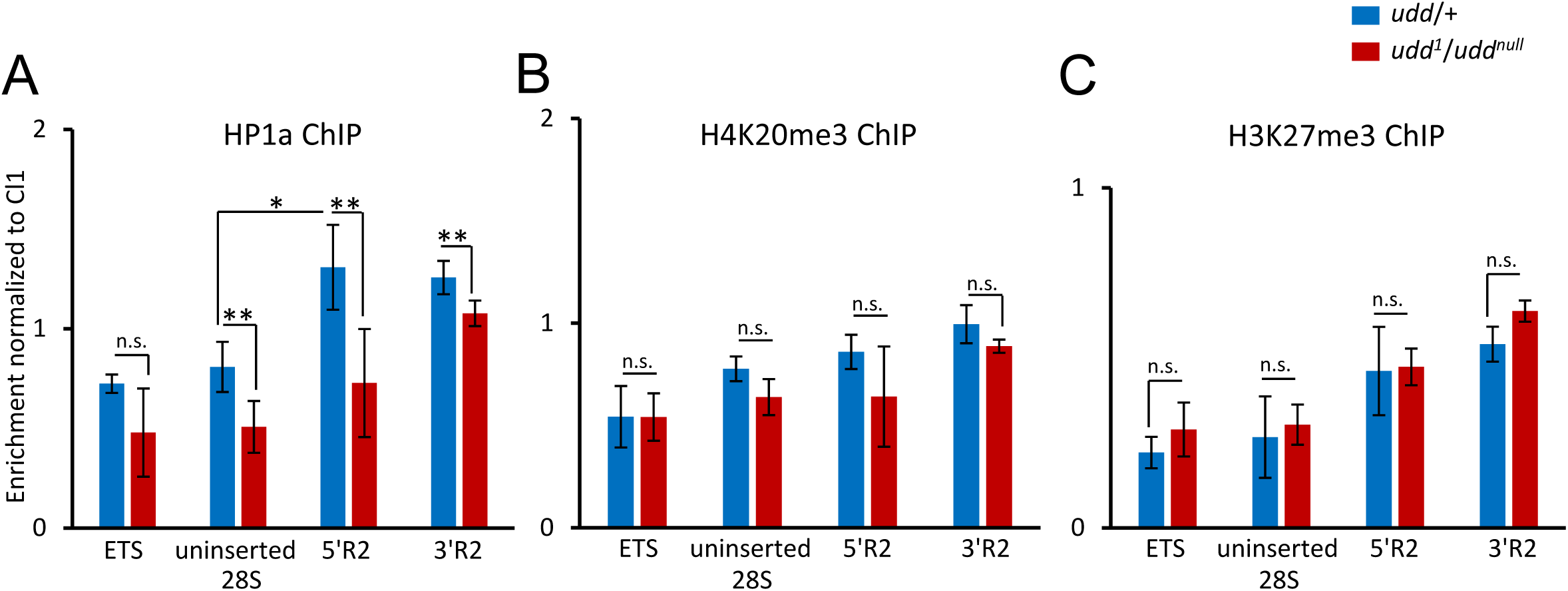
The effect of Udd loss on repressive chromatin marks at rDNA. ChIP-qPCR analysis of HP1a (**A**), H4K20me3 (**B**) and H3K27me3 (**C**) levels in *udd*^*1*/^*udd*^*null*^ (red bars) and *udd*/+ (blue bars) ovaries. p-values calculated using Student’s t-test, * p<0.003, ** p<0.01, n.s. = not significant.

